# Visual-olfactory integration in the human disease vector mosquito, *Aedes aegypti*

**DOI:** 10.1101/512996

**Authors:** Clément Vinauger, Floris Van Breugel, Lauren T. Locke, Kennedy K.S. Tobin, Michael H. Dickinson, Adrienne Fairhall, Omar S. Akbari, Jeffrey A. Riffell

## Abstract

Mosquitoes rely on the integration of multiple sensory cues, including olfactory, visual, and thermal stimuli, to detect, identify and locate their hosts [1–4]. Although we increasingly know more about the role of chemosensory behaviours in mediating mosquito-host interactions [1], the role of visual cues remains comparatively less studied [3], and how the combination of olfactory and visual information is integrated in the mosquito brain remains unknown. In the present study, we used a tethered-flight LED arena, which allowed for quantitative control over the stimuli, to show that CO_2_ exposure affects target-tracking responses, but not responses to large-field visual stimuli. In addition, we show that CO_2_ modulates behavioural responses to visual objects in a time-dependent manner. To gain insight into the neural basis of this olfactory and visual coupling, we conducted two-photon microscopy experiments in a new GCaMP6s-expressing mosquito line. Imaging revealed that the majority of ROIs in the lobula region of the optic lobe exhibited strong responses to small-field stimuli, but showed little response to a large-field stimulus. Approximately 20% of the neurons we imaged were modulated when an attractive odour preceded the visual stimulus; these same neurons also elicited a small response when the odour was presented alone. By contrast, imaging in the antennal lobe revealed no modulation when visual stimuli were presented before or after the olfactory stimulus. Together, our results are the first to reveal the dynamics of olfactory modulation in visually evoked behaviours of mosquitoes, and suggest that coupling between these sensory systems is asymmetrical and time-dependent.

## Results and Discussion

To detect and locate suitable hosts, mosquitoes rely on multiple sensory cues, including olfactory, visual, and thermosensory information [1–5] while flying through a dynamic environment [6]. Whereas responses to olfactory (for review [7–9]) and, to a lesser extent, thermal stimuli [10,11] have been well studied, comparatively less is known about how visual cues evoke behavioural responses in mosquitoes (but see [12,13]) and how olfaction and vision are integrated. A recent study showed that CO_2_ detection activates a strong attraction to visual features that is critical for mediating interaction with close-range cues like heat and other host volatiles [3]. However, because this study relied on free-flight assays, it was difficult to control the sensory experience of the mosquitoes, which is a function of both their trajectories through space and the spatiotemporal distribution of the stimuli within the wind tunnel. In other animals, ranging from bees to humans, prior exposure to a visual stimulus can modify olfactory responses [14-16], and *vice versa* [17]. It thus remains an open question how the temporal order and interval between detection of olfactory and visual stimuli influences mosquito behaviour and processing in the brain.

In contrast to free-flight assays, tethered flight experiments [18] offer the means to manipulate the temporal and quantitative nature of multimodal stimulation (*e.g*., [19]) and renders possible a behavioural analysis at the scale of wingbeat kinematics [18]. In mosquitoes, tethered flight has been employed to characterize responses to both visual [13] and olfactory cues [20]. Here, to study the integration of multimodal host signals in *Ae. aegypti*, we placed tethered mated females within an LED cylindrical arena [18] that permitted simultaneous presentation of olfactory stimuli with the motion of high-contrast visual objects. Within the arena, a mosquito was centred between an air inlet and a vacuum line aligned 30° diagonally from the vertical axis (Fig. 1A). The air inlet was positioned 12 mm in front of, and slightly above, the mosquito’s head to target the antennae from an angle of 15°. We monitored the mosquitoes’ responses to visual and olfactory cues by tracking changes in wingbeat frequency and amplitude. These measures are proxies for changes in the flight speed of the insect, as increases in these signals are indicators of the surge response elicited by the detection of an odour plume in free-flight [21–25]. The turning tendency, defined as the difference between the left and right amplitudes, was used to monitor directional changes attempted by the mosquitoes. These turn attempts were used as a proxy for determining the aversive or attractive nature of the visual stimuli.

**Figure 1.**
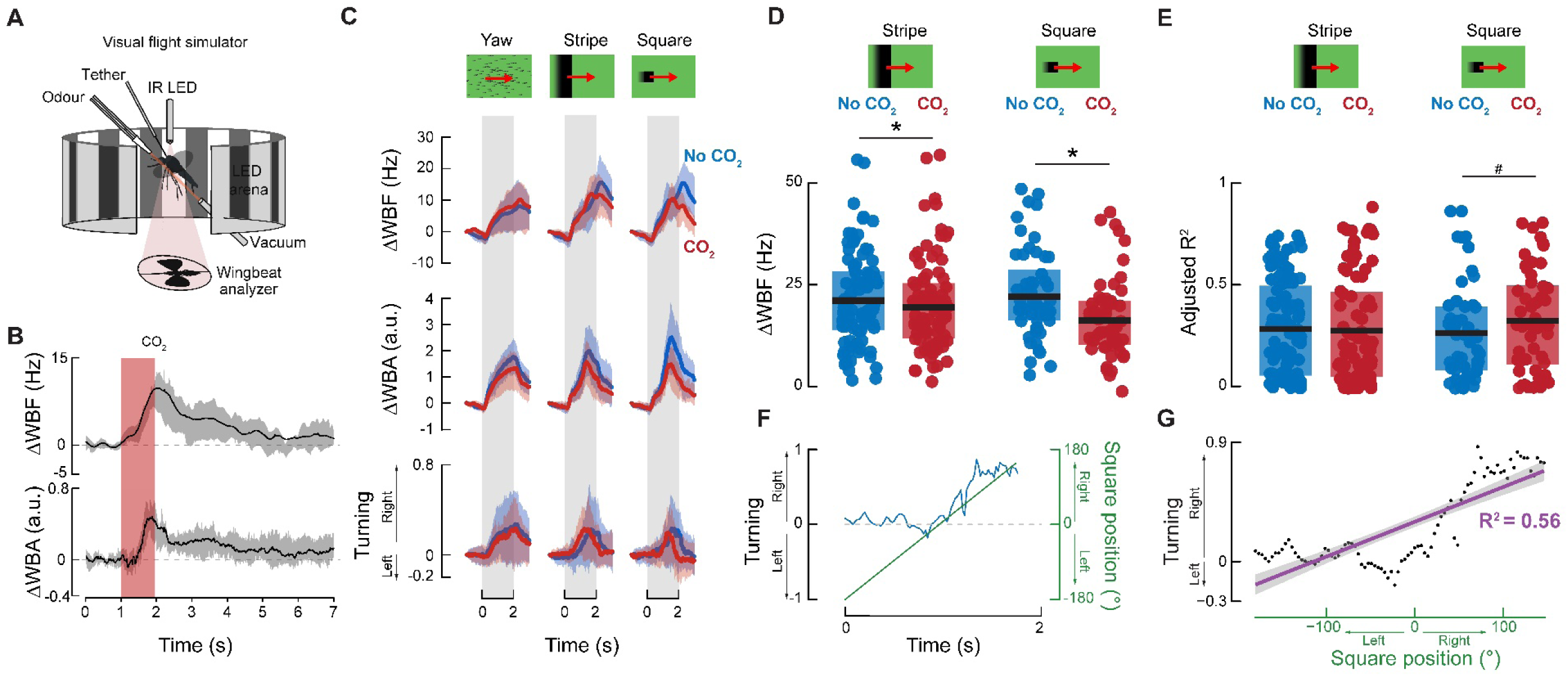
CO_2_ modulates mosquitoes’ responses to small field rotating visual objects. **(A)** Visual flight simulator (adapted from [14,15]) used to record wing kinematics from a tethered mosquito. **(B)** Stimulus**-**trigger-averaged changes in wingbeat frequency (ΔWBF) and amplitude (ΔWBA) (solid lines) in response to a 1-sec pulse of 5% CO_2_ (red bar). Shaded areas represent the mean ± the first quartiles. **(C)** Mean responses of mosquitoes to a panel of visual stimuli. *Top*: normalized wingbeat frequency, *middle*: amplitude and *bottom*: turning changes induced by the visual stimuli. Plotted are the mean responses to visual stimuli in the absence (blue lines) and presence (red lines) of CO_2_. Shaded areas denote the first and last quartiles around the mean. **(D)** Mean (±first and third quartiles) wingbeat frequency responses to stripes and squares with (red dots) and without (blue dots) CO_2_. Asterisks denote significant differences between CO_2_ and no CO_2_ treatments (p < 0.05; n = 51-116). **(E)** Mean (±first and third quartiles) adjusted R^2^ values of fitted linear model defined as: mosquito.turning = α × object.position + β × object.velocity. Responses to stripes and squares with (red dots) and without (blue dots) CO_2_. (# indicates a p-value of 0.09; n = 51-116). **(F)** Representative example of a single mosquito’s turning response (blue line) relative to the position of a moving square (green line) in the absence of CO_2_. **(G)** Example fitted linear model (purple line) illustrating how the turning direction of the mosquito depicted in (F) is explained by the position of the square.

### Tethered mosquitoes respond to pulses of CO_2_ by increasing their wingbeat frequency

To first characterize the behavioural responses of tethered mosquitoes to host-emitted cues, the mosquitoes were maintained in a static (*i.e*., open-loop and immobile) visual environment composed of multiple dark vertical stripes (22.5° wide x 60° tall, spaced by 22.5°) and presented with pulses of CO_2_, a strong attractant that is considered to be the most important sensory cue used by mosquitoes to find their hosts [26]. The concentration of CO_2_ emitted by humans is ~4.5%, and mosquitoes typically fly upwind in filamentous plumes of CO_2_, within which the concentrations falls to background levels (0.03-0.04%) as the distance from the host increases [6,27-29]. We therefore tested mosquito responses to CO_2_ at different concentrations (1 to 10%) and with pulses of various durations (0.5 to 20 sec) to simulate the temporal statistics of a natural plume.

To characterize the change in flight dynamics upon olfactory and visual stimulation, we quantified the change in wingbeat frequency (ΔWBF) and wing-stroke amplitude (ΔWBA) (Fig. S1). Increases in WBF and WBA are correlated with the upwind surge that mosquitoes exhibit in free flight. In our tethered preparation, we found increases in both parameters when the animals were presented with CO_2_, but found more consistent changes in the WBF (Fig. S1A). In the absence of olfactory stimuli, tethered mosquitoes exhibited a stable wingbeat frequency that ranged from 397 to 579 Hz. Short, 1 sec pulses of 5% and 10% CO_2_ elicited transient increases in wingbeat frequency above this baseline (9 ± 3 Hz, and 4 ± 1 Hz respectively; *t*-tests, *t*_14_=2.82, p=0.01 and *t*_15_=3.2, p=0.005, respectively) (Figs. 1B, S1). Pulses of 1% and 2.5% CO_2_ did not elicit significant changes in wingbeat frequency and were similar to control pulses of nitrogen (N_2_) (Fig. S1C; *t*-tests, p>0.05). The duration of the CO_2_ pulse also had a strong influence on the time course of wingbeat responses, with longer pulse durations – particularly for 5% and 10% concentrations – causing a significantly longer decay time back to baseline (Table S1).

### Carbon dioxide modulates responses to certain visual stimuli

Given the robust responses obtained with 1 sec pulses at 5% CO_2_, which is close to human emissions [28], we chose this concentration and pulse duration to investigate the effect of CO_2_ on the responses to visual stimuli. Previous seminal studies by Kennedy and others have examined the visual responses of mosquitoes in both free and tethered flight preparations and established the attraction of mosquitoes to dark objects [13, 31–33]. A more recent study [3] showed that female *Ae. aegypti* mosquitoes are attracted to dark visual objects only after encountering a plume of CO_2_. In these experiments, the visual object was located well outside the CO_2_ plume, suggesting that simultaneous experience of the two cues is not necessary; rather, CO_2_ causes a change in visual response that persists after odour encounter.

To characterize the effect of CO_2_ on tethered mosquitoes’ visual responses, we used a panel of visual stimuli in the absence of odour, including small and large looming objects, drifting objects, and large-field patterns of optic flow [34] (Fig. 1, S2). In the absence of CO_2_, different open-loop visual stimuli triggered different responses. For example, multiple vertical stripes or starfield patterns creating progressive optic flow (i.e., expanding symmetrically from the centre of the arena), and a single square or bar performing one rotation around the mosquito elicited the strongest increases in wingbeat frequency (ΔWBF) (Fig. S2), whereas regressive motion patterns (either stripes or starfields) elicited only moderate responses (Fig. S2). We also observed that mosquitoes responded with a larger increase in wingbeat frequency to looming stimuli (i.e. a square growing from 3.75° [1 pixel] to 60° wide [16×16 pixels]) than to contracting stimuli (i.e., a square shrinking from 60° to 3.75° wide).

To simulate a plume encounter, visual stimuli were preceded by a 1 sec pulse of 5% CO_2_. The CO_2_ stimulus had a small but significant effect on their responses towards discrete objects such as stripes and squares (paired *t-*test: *t*_51-85_ > 1.75, p < 0.05) (Fig. 1C-D), but not wide field stimuli (paired *t*-tests: *t*_69-116_<3.001, p>0.05). For horizontally drifting visual patterns (*e.g.*, starfield yaw, stripe, square), the mosquitoes followed the motion direction (Fig. 1C). To further quantify the extent to which mosquitoes tracked the object during its presentation (Fig. 1F), we fitted a linear model based on the visual object position and the mosquito’s turning response (mosquito.turning = α × object.position + β × object.velocity; Fig. 1G). The adjusted coefficient of correlation R^2^ was used as a metric of how well the turning of the mosquito is explained by the motion of the object. For squares, although not significant, the presence of CO_2_ tended to increase this relationship between object motion and the turning response of the mosquito (*t*-test: *t*_100_=-1.2908, p=0.09) (Fig. 1E). It is worth noting that mosquitoes exhibited this strong turning tendency towards both stripes and squares and in both the presence and absence of odour (Fig. 1C). These results diverge from results from other dipterans such as *D. melanogaster* which turn towards stripes taller than ~25° and turn away from smaller objects (8-25° in height) in the absence of odour [35].

### CO_2_-induced modulation of visually mediated behaviour is time-dependent

Previous free-flight experiments in mosquitoes have shown that attraction to visual objects is odour-mediated and can last on the order of seconds [3]. These experiments, however, were unable to control the time interval between the olfactory and visual cues or the effect of the size of the visual objects. To better understand these effects, we presented mosquitoes with moving squares of different sizes (3.75-22.5°) preceded by either no CO_2_, or a 1 sec pulse of CO_2_ delivered 1, 5, or 10 seconds before the onset of the visual stimulus (Fig. 2A). For the larger squares (7.5° to 22.5°), mosquitoes exhibited strong turning responses towards the visual object (Fitted Linear Model: *F*_1,1248_ > 12.23, p<0.05; Fig. 2D) that were not significantly different between time periods after CO_2_ stimulation (p>0.05). However, for the smallest square, CO_2_ had a significant impact on the turning responses, with the strongest levels of correlation between the response and object position occurring at 5 and 10 seconds after CO_2_ presentation (*t-*test; *t*_50_=- 3.536, p<0. 0.001). In addition to changes in turning responses, CO_2_ modulated the mosquito’s wingbeat frequency: CO_2_ delivered 1 sec before the visual stimulus (22.5° square) elicited a significant reduction in the wingbeat frequency response to the moving square (paired *t*-test: *t*_18_=-1.72, p = 0.046). Across stimuli and intervals, the change in wingbeat frequency was significantly influenced by the inter-stimulus timing (ANOVA: *F_3,294_=4.53*, p<0.01)(Fig. 2; Table S2), and the size of the square (p<0.001)(Fig. 2B). For instance, the bigger the square, the larger the response, and there was a strong time-dependency in these effects occurring 1 sec after odour exposure (Tukey post-hoc test: p<0.01). This time-dependency was consistent between squares of different sizes, where all but the smallest square (3.75°) showed a mean decrease at 1 sec (Fig. 2B). Together, these results suggest that CO_2_ suppresses turns from the visual targets (typically associated with an increase in WBF) and demonstrate that prior stimulation with odour influences visual object tracking in a time-dependent manner.

**Figure 2.**
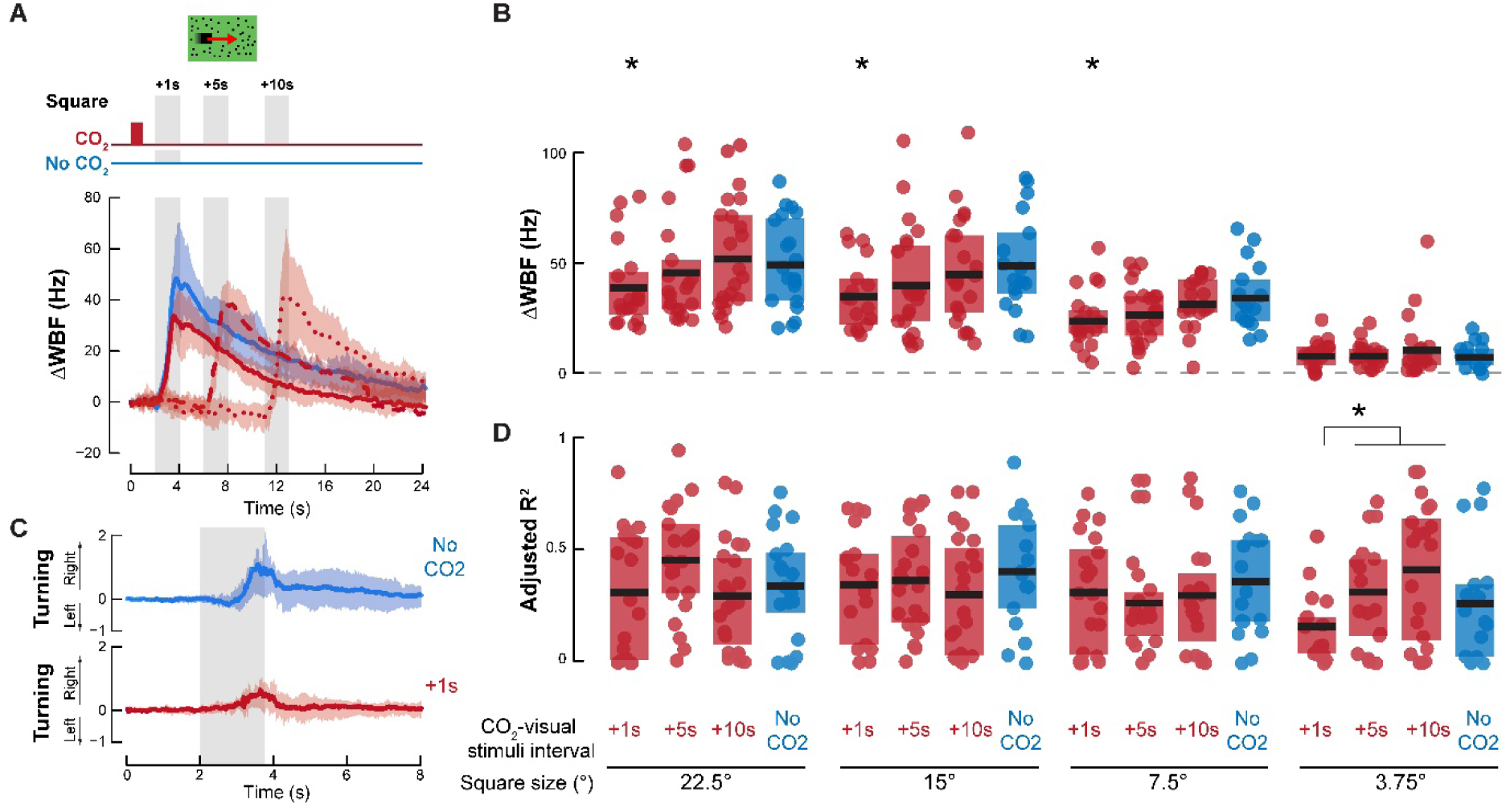
The effect of CO_2_ peaks at 1 sec and is a function of the spatial characteristics of the visual object. **(A**) *Top*: Schematic of stimulus time series; and *bottom*: Stimulus-triggered average wingbeat responses to a moving (yaw) square without CO_2_ (blue line) or 1, 5 or 10 sec after stimulation with CO_2_ (red lines). Shaded areas denote the first and last quartiles around the mean (n=13 mosquitoes). **(B)** The change in wingbeat frequency for different square sizes and intervals between CO_2_ and visual stimuli. Both the interval between stimuli, and the square size, significantly modified the mosquito’s responses (two-way ANOVA: p < 0.05). Bars are the means (±first and third quartiles); asterisks denote significant differences from the “No CO_2_” treatment (p < 0.05). **(C)** Mean (±first and third quartiles) turning response of mosquitoes to a moving 22.5° wide square in the absence (top, blue) or presence of CO2, delivered 1s prior to visual stimulation (bottom, red). Grey shaded area indicates the duration of visual stimulation. **(D)** Mean (±first and third quartiles) adjusted R^2^ values of fitted linear model of turning responses defined as: mosquito.turning = α × object.position + β × object.velocity, for different square sizes and intervals between CO_2_ and visual stimuli. Asterisks denote significant differences (p < 0.05).

It is difficult to directly compare the behavioural responses of tethered and freely flying insects because the tethered animals lack proper proprioceptive and haltere feedback, and the visual contrast in the LED arenas is often greater than what they experience in a natural or wind tunnel setting. In our tethered preparation, instead of CO_2_ gating their attraction to an immobile square stimulus as it was observed in free flight [3], CO_2_ increased the animals’ tracking fidelity of moving objects. This modulation is consistent with, yet more subtle than, their free flight behaviour. It is worth noting that the difference between free flight [3] and tethered behaviour is also less pronounced in *Drosophila* [36]. Although the behavioural modulation we observed was subtle, quantifying the modulation allowed us to design a tethered preparation in which we could use calcium imaging to observe the neural basis for the behavioural modulation.

### Odour selectively modulates optic lobe responses

Given the modified behavioural responses to visual stimuli after odour stimulation, we took the first steps towards understanding where in the brain olfactory and visual information is integrated by monitoring neural activity in both the antennal lobe, primary processing centre of olfactory information, and the lobula, a region in the mosquito optic lobe. We imaged calcium levels in groups of neurons with two-photon excitation microscopy (Figs. 3A, 4A,H) in an *Ae. aegypti* line genetically modified to express a genetically encoded Ca^2+^ indicator driven by a ubiquitin promotor (termed *PUb*-*GCaMP6s*). Although this mosquito line does not permit cell-type specific expression and labelling, *PUb*-*GCaMP6s* is expressed strongly in axons and neuropil in the central nervous system and shows clear stimulus-evoked responses of specific neuronal types in the visual and olfactory brain regions (*e.g.*, antennal lobe projection neurons) (Figs. 3,4) [37].

**Figure 3.**
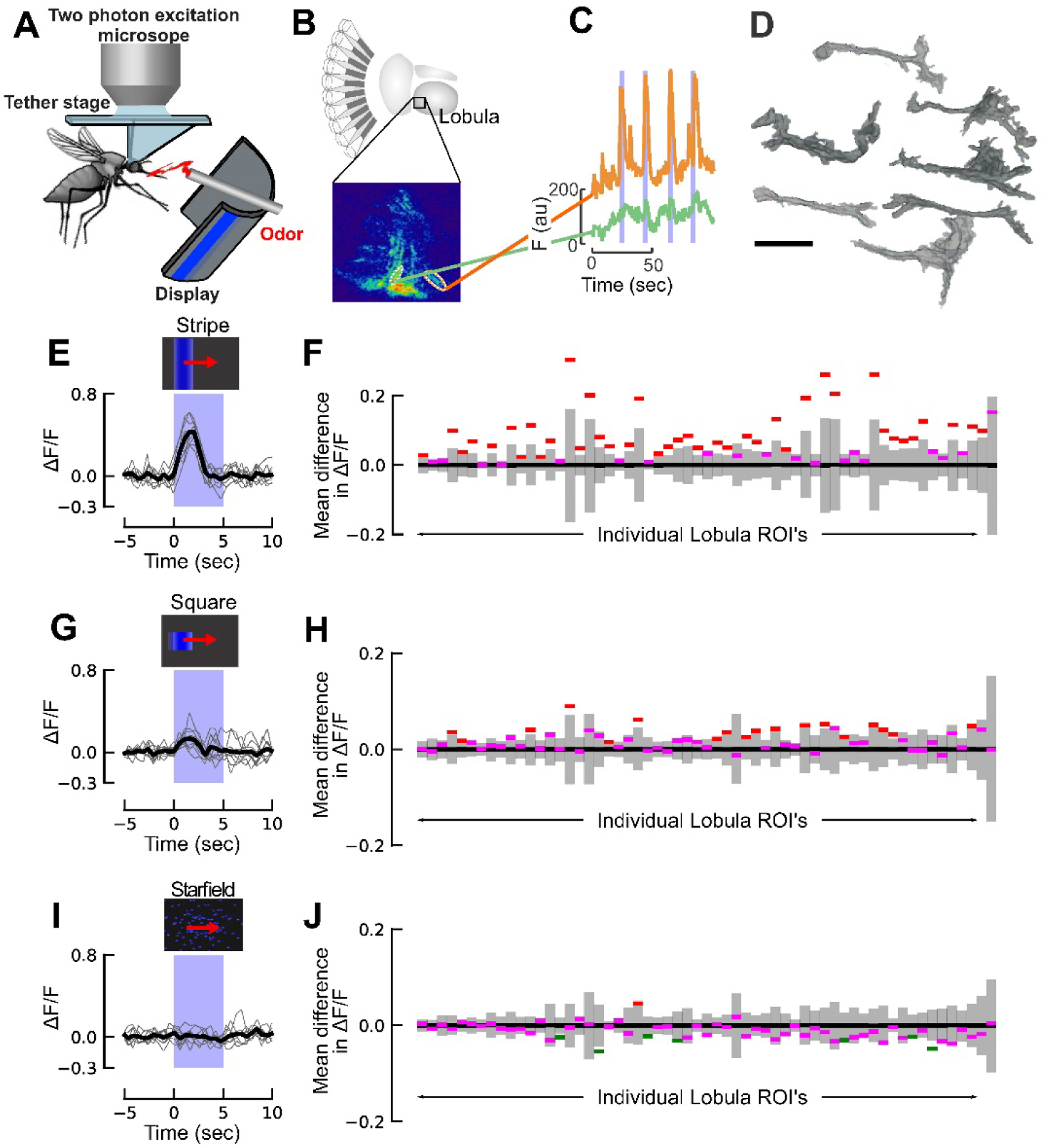
Lobula responses to visual stimuli. **(A)** Schematic of the two-photon setup used to record calcium dynamics in the mosquito antennal and optic lobes. **(B)** Diagram of the *Ae. aegypti* optic lobe, highlighting the lobula and a representative pseudocoloured image of the scanning plane. **(C)** Representative time traces of two lobula ROIs showing different levels of stimulus-evoked responses (orange and green) to the visual stimulus (blue bars). **(D)** Representative 3-D reconstructions of ROIs that showed evoked responses to visual stimuli. Certain ROIs (right) showed tree-like dendritic branching, whereas others showed more columnar morphology (left). Although we were unable to assign imaged neuropil to orthologous neurons in other dipteran species, like *D. melanogaster*, intriguing similarities may exist based on their neuroanatomy, such as the LC or LT cells [53]. Scale bar: 20 µm. **(E)** ∆F/F time trace for ROI# 16, showing the strong response to the bar stimulus. **(F)** Two thirds of the ROI’s we imaged in the lobula significantly responded to motion of a moving bar. Red lines represent the responses of ROIs that are significantly greater (p<0.05) than the null distribution (grey bars); purple lines represent those ROI responses that are not significantly different from the null (grey bars) distribution (p>0.05). To analyze the significance of the change in fluorescence of each ROI in response to a moving bar, we compared the mean fluorescence during the first two seconds of visual motion to the mean fluorescence for a two second period two seconds prior to when the visual stimulus began. For both of these analyses, we calculated ∆F/F relative to the 5 seconds prior to the time of interest, which controls for slow changes in the fluorescence signal. The results are plotted as in Fig. 4, and the ROI’s in this figure are sorted in the same order shown in Fig. 4, allowing for a direct comparison. **(G)** Same ROI as E, but in response to a moving square. **(H)** Same as F, but for a moving square. Twenty-eight percent of the ROIs showed significant responses to the moving square (p<0.05). **(I)** Same ROI as E, but in response to a moving star field. **(J)** Same as F, but for a moving star field. Only 13% of the ROIs showed significant responses to the moving square (p<0.05). Green lines represent the responses of ROIs that are significantly less (p<0.05) than the null distribution (grey bars).

**Figure 4.**
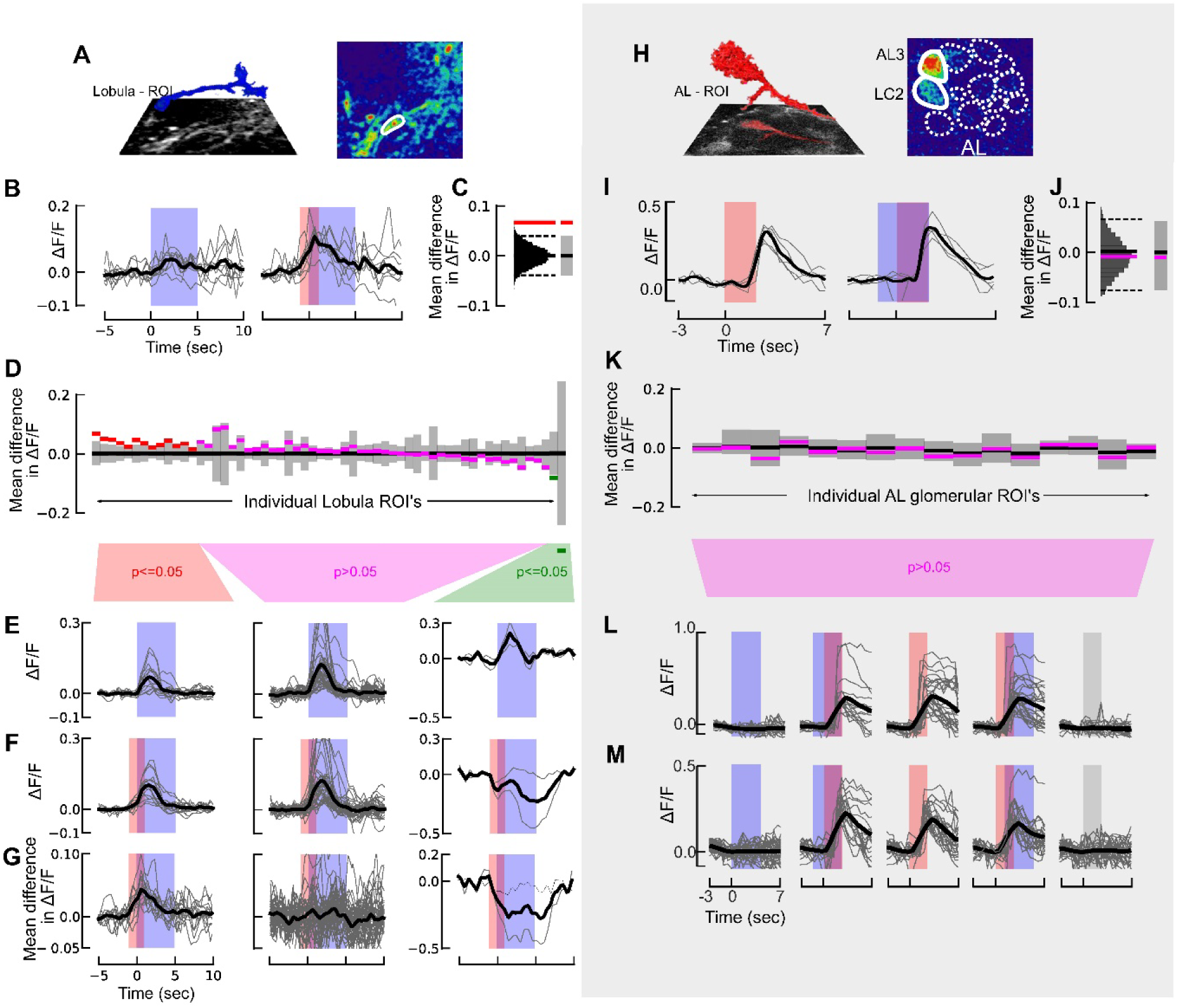
Calcium imaging of visual responses in the mosquito antennal and optic lobes reveals asymmetric neuromodulatory effect of odour. **(A)** 3D reconstruction of a lobula ROI inset above the imaging plane (left). (Right) pseudocolour plot of the calcium fluorescence during the presentation of a visual stimulus. **(B)** Time series of ∆F/F in one ROI for 9 presentations of the visual stimulus (bar) without an odour stimulus (left) and with an odour stimulus preceding the visual stimulus (right). **(C)** To assess the difference in the response for the odour and no-odour experiments, we calculated the difference in the mean ∆F/F during the visual stimulus period for the two experiments (red line). We then generated a null distribution by pooling the data from both experiments and bootstrapping 10,000 pairwise differences from this combined dataset (gray histogram). If the actual difference lies outside of 95% of this bootstrapped distribution (dashed lines), the difference is significant (p<=0.05; depicted by red line). **(D)** As in C, but for the 59 lobula ROIs where the difference in mean ∆F/F for the odour and no-odour case, calculated as in C. **(E)** ∆F/F time traces for the “no odour” visual stimulus, split into the three statistical groups shown in D. Thin traces show the average response for each ROI across 9 trials. **(F)** ∆F/F time trace for odour+visual stimulus experiments, as in E. **(G)** Difference in the mean ∆F/F time traces shown in E and F for each ROI. **(H)** 3D reconstruction of the AL3 projection neuron (red) above the imaging plane (left), and (right) pseudocolour plot of the *Ae. aegypti* AL at the 30µm imaging depth. AL glomeruli are depicted by dashed lines; the nonanal-responsive AL3 and LC2 glomeruli are depicted by the white solid lines. **(I)** Time series of ∆F/F in one ROI for 5 odour stimulations without a visual stimulus (*left*) and with a visual stimulus preceding the odour stimulus (*right*). **(J)** As in C, but for the AL3 glomerulus in I. **(K)** Difference in mean ∆F/F for the odour and no-odour case for each of the 16 glomerular ROI’s, calculated as in C. **(L)** ∆F/F time traces for the AL3 glomerulus in the different treatments: visual presentation alone; visual presentation preceding the odour; odour alone; odour stimulus preceding the visual presentation; and no odour (mineral oil) control. Thick trace (black) is the mean from 8 mosquitoes; thin grey traces are the individual stimulations across all animals (n=4 or 5 per mosquito). **(M)** As in L, but for the LC2 glomerulus.

We first focused on the lobula, as this area is comprised of target-selective neurons in other dipteran species [38–41]. Using the *PUb*-*GCaMP6s Ae. aegypti* line coupled with two-photon excitation microscopy, we imaged dendrites and axons at approximately 30-100 μm depth from the ventral surface of the lobula, as neurons in this region showed strong GCaMP6s expression and exhibited unambiguous responses to visual and olfactory stimuli (Fig. 3B,C; Movie S1). ROIs were manually selected based on the following criteria: strong GCaMP6s expression in dendrites and axons (1.02 to 15.9-times background levels), surface areas of 40-100 μm^2^, and responses to visual or olfactory stimuli in at least one of the ROIs in the imaging plane. Moreover, the strong GCaMP6s expression permitted 3D reconstructions of the imaged neuropil (Figs. 3D; 4A,H). As a first step to characterizing lobula responses, we presented the mosquito a single moving stripe, square (15°), or star-field pattern. We imaged 59 ROIs across 6 individual mosquitoes, and each stimulus type was presented 9 times. For each ROI, we compared the mean fluorescence during the first two seconds of visual motion to the mean fluorescence two seconds prior to when the visual stimulus began, and compared these two datasets to a null distribution of 10,000 bootstrapped pairwise differences drawn from the combined datasets. Results from these analyses revealed that the moving stripe evoked strong and significant responses in approximately 67% of the ROIs (Fig. 3E,F), while presentation of the square evoked weaker responses in the population with approximately 28% of the ROIs showing responses (Fig. 3G,H). In contrast, few ROIs exhibited significant responses to wide-field motion of the star-field (13% responding), with the majority of these ROIs showing significant suppression, rather than strong excitation evoked by the bar and square (Fig. 3I,J).

To examine how odour modulates visually-evoked responses in these ROIs (Fig. 4A), we presented the mosquito with a moving stripe, with and without CO_2_ pulses prior to the onset of the visual stimulus (Fig. 4A,B). Similar to the above analysis, for each ROI we assessed the difference between the odour and no-odour responses by comparing the differences using a bootstrap analysis of the combined odour and no-odour datasets (Fig. 4B,C). In 13 of these ROIs, the visually evoked responses were significantly larger (p<0.05) when preceded by CO_2_, and in 2 of them, the responses were significantly smaller (p<0.05) (Fig. 4D-G). Note that at a cut-off of p=0.05, we would only expect 3 ROIs to exhibit a significant difference by chance, suggesting that most of the 15 ROIs that exhibited different visual responses are indeed being modulated by the preceding odour. Next, we took a closer look at the 13 ROIs that exhibited modulation. These ROIs included data from 5 of the 6 individuals. In both the modulated and unmodulated ROIs we found a strong correlation between the responses to the visual stimulus and the odour + visual stimulus (Linear regression for modulated ROIs: R^2^=0.89; p<0.0001 and unmodulated ROIs: R^2^=0.89; p<0.0001). Thus, the average response for the positively modulated ROIs was similar with and without odour presentation, but for each individual ROI, the preceding odour increased the response to the visual stimulus by 0.037 (47%) (Fig. 4F,G).

Next, we tested whether the modulated ROIs might respond to odours in the absence of any visual stimuli. We found that on average these ROIs responded to a 2-second pulse of odour with an increase in fluorescence of 0.015, which explains 40% of the visual modulation, suggesting a non-additive modulatory effect (Fig. S3A). ROIs that did not exhibit modulation showed no response to the odour stimulation, and ROIs that exhibited negative modulation also exhibited reduced fluorescence in response to the odour pulse (Fig. S3A). Previous research has shown that optic lobe neurons may be sensitive to mechanosensory input [42,43]. We therefore tested elevated pulses of airflow approximate to those used in our olfactory experiments. Results showed no significant increase in calcium dynamics corresponding to the timing of the air pulse (Fig. S3B), suggesting that the mechanical aspect of the stimulus did not lead to modulation in the lobula. However, more carefully targeted experiments will be needed to tease apart how odour and mechanical stimuli work together to modulate neuronal responses in the lobula.

### Visual stimuli do not modulate responses in the mosquito antennal lobe

We next examined whether visual stimuli modulated responses in the antennal lobe (AL), as this brain region provides neuroanatomically identifiable glomeruli tuned to different odorants and receives feedback from higher-order regions that also receive visual input [44]. Calcium imaging experiments were conducted with eight animals using stimuli similar to those described in the previous section, in which an individual mosquito was presented with visual stimuli (moving stripe) with and without pulses of CO_2_ and nonanal, another host-emitted odorant [45]. Prior to odour stimulation, glomerular boundaries were discernible based on the baseline GCaMP6s expression, although significant calcium transients were not apparent. However, upon odour stimulation the ventral AL glomeruli (AL3 and LC2), which are responsive to nonanal and other host odours, showed strong calcium responses that were time-locked to the odour stimulus (Kruskal-Wallis test: χ^2^ >43.7, p<0.0001; multiple comparisons relative to control: p<0.001) (Fig. 4H,I). When stimulated, dendritic arbours filling the AL3 and LC2 glomeruli and axons projecting into the coarse neuropil became observable (Fig. 4H). When visual stimuli were presented with the odour – either 1 sec before, or 1 sec after –, there was no difference in response compared to when odour was delivered alone (p>0.99) (Fig. 4I-M). Moreover, responses to isolated visual stimuli were not significantly different from the mineral oil (no odour) control (p>0.97) (Fig. 4L,M), demonstrating that the observed wingstroke responses to coupled olfactory and visual stimuli were not correlated with modulated responses in the AL.

## Conclusions

As mosquitoes navigate to human hosts, they will experience an odour plume that varies in time and intensity, and a visual target that varies in angular size [4,32,33]. To simulate these conditions in the laboratory, we used a flight arena that permits the fine-scale control of the size of visual object – corresponding to a human that is 1 to 20 m away from the mosquito [32,33] – and odour concentrations and durations typical to what a mosquito experiences when encountering a plume [6,28]. Results from these experiments demonstrate that olfactory input modulates subsequent responses by *Ae. aegypti* mosquitoes to dark, moving, smaller-field visual stimuli, possibly reflecting finer flight kinematic control towards visual objects indicative of blood hosts. Furthermore, our results show that modulation of visually evoked behaviours is limited to less than 10 sec after odour stimulation. These behavioural responses are reflected in the functional calcium imaging of neuropil in the lobula, where 20% of the ROIs showed an increase or decrease in their fluorescence when the odour preceded the visual stimulus. Although we did see modulation of several ROIs in the lobula, not all exhibited strong changes in their responses when odour and visual stimuli were presented together. By contrast, in free-flight experiments, odour encounters qualitatively gated attraction to visual objects. Our imaging results suggest that this free flight behaviour is the result of two levels of modulation: odour-induced modulation occurring in or upstream of the lobula, and additional modulation in other downstream brain-regions involved in processing visual information [34] that we have not yet recorded. In contrast to the odour-induced modulation of visual responses, we did not observe visual-induced modulation of olfactory responses in the antennal lobe, suggesting that any such modulation, if it occurs, must happen downstream of the antennal lobe, perhaps in the higher-order brain regions that receive both olfactory and visual inputs [44-48].

Research in a variety of animals has shown that olfactory and visual stimuli can symmetrically, or asymmetrically, modulate each other depending upon the context. In humans, when visual and olfactory cues conflict with each other, vision is the dominant modality and overrides olfaction [2]. A classic example comes from unfortunate University of Bordeaux oenology students who tested “red” wines that were in fact white wines with red food colouring; students used terminology and descriptors typically reserved for red wine and avoided those for white [49]. However, prior exposure to an odour can also facilitate responses to visual stimuli, especially when the cues are congruent; for instance, the scent of a rose has been shown to facilitate the identification of a picture of a rose [17]. Studies have shown that visual and olfactory integration and modulation occurs in the higher-order areas of the brain, like the hippocampus and orbitofrontal cortex, rather than in the sensory areas (e.g., olfactory bulb or dorsal lateral geniculate nucleus) [14].

Behavioural experiments with mosquitoes and other insects have shown that they too integrate multiple sensory modalities to function efficiently and robustly in complex environments [3,4,21,43]. Recent advances in genetic tools have allowed scientists to begin probing where in the brain this integration occurs. Like mammals, insects exhibit symmetric sensory integration and modulation in higher order brain areas, such as the mushroom bodies [47,48,50] and central complex [51]. However, in contrast to mammals, insects also exhibit sensory integration and modulation in early sensory areas, such as in the antennal lobe [52] and optic lobe [42,43]. Nonetheless, the degree to which sensory integration occurs in more peripheral processing areas for insects remains an open question. Our experiments suggest that in the mosquito sensory modulation is asymmetric: odour modulates vision, but not vice versa. Making a direct comparison between the two systems is difficult, as there are more regions and connections in the visual system compared to the olfactory system [53-55]. Prior studies, however, have suggested that an area just downstream from the lobula, referred to as the optic glomeruli, shares many anatomical similarities with the antennal lobe [54], making these brain regions ideal for functional comparisons. Although we did not image from the optic glomeruli, the fact that we saw olfactory modulation in the lobula, one step before the optic glomeruli, but no visual modulation in the antennal lobe, suggests that the modulation is indeed asymmetric in these loci in the brain.

Why might sensory modulation in the mosquito be asymmetric? Insects such as mosquitoes have relatively poor visual resolution (5°, compared to humans’ 0.02°). Thus, for a mosquito, vision is unlikely to provide information about what something is. Instead, the odour may provide information for what the animal is smelling, while vision provides information for where the odour is located. These differences might explain the asymmetric sensory modulation we observed. Future comparative studies across species with varying degrees of resolution in sensory modalities are needed to address this hypothesis. Finally, there is a growing understanding of the molecular and neurophysiological bases of olfactory behaviours in mosquitoes [9], but we know comparatively little about their visual behaviours despite its importance for locating hosts and selection of biting sites [3, 56, 57]. Our results here provide motivation for addressing this research gap, as well as identifying the mechanisms by which olfactory input modulates other sensory systems. Fortunately, thanks to the recent development of new genetic tools, these types of experiments are now possible.

## Acknowledgements

We thank B. Nguyen for mosquito colony maintenance, J. Tuthill and P. Weir for comments and help with the arena experiments, G. Wolff for comments and imaging assistance, and D. San Alberto for technical support. We acknowledge the support of the Air Force Office of Sponsored Research under grant FA9550-14-1-0398 and FA9550-16-1-0167, National Institutes of Health under grant NIH1RO1DCO13693-04, an Endowed Professorship for Excellence in Biology (J.A.R.), the University of Washington Innovation Award. O.S.A was supported in part by NIH grants 5K22AI113060 and 1R21AI123937.

## Author Contributions

C.V., F.V.B., A.F., M.H.D. and J.A.R. conceived the study. C.V., L.T.L. and K.S.T. participated in the execution and analysis of the arena assays. O.S.A. generated the GCaMP6 mosquitoes. J.A.R. conducted the imaging assays and C.V., F.V.B. and J.A.R. analysed the imaging data. C.V., F.V.B. and J.A.R. wrote the paper, and all authors edited the manuscript.

---

**Figure S1.**
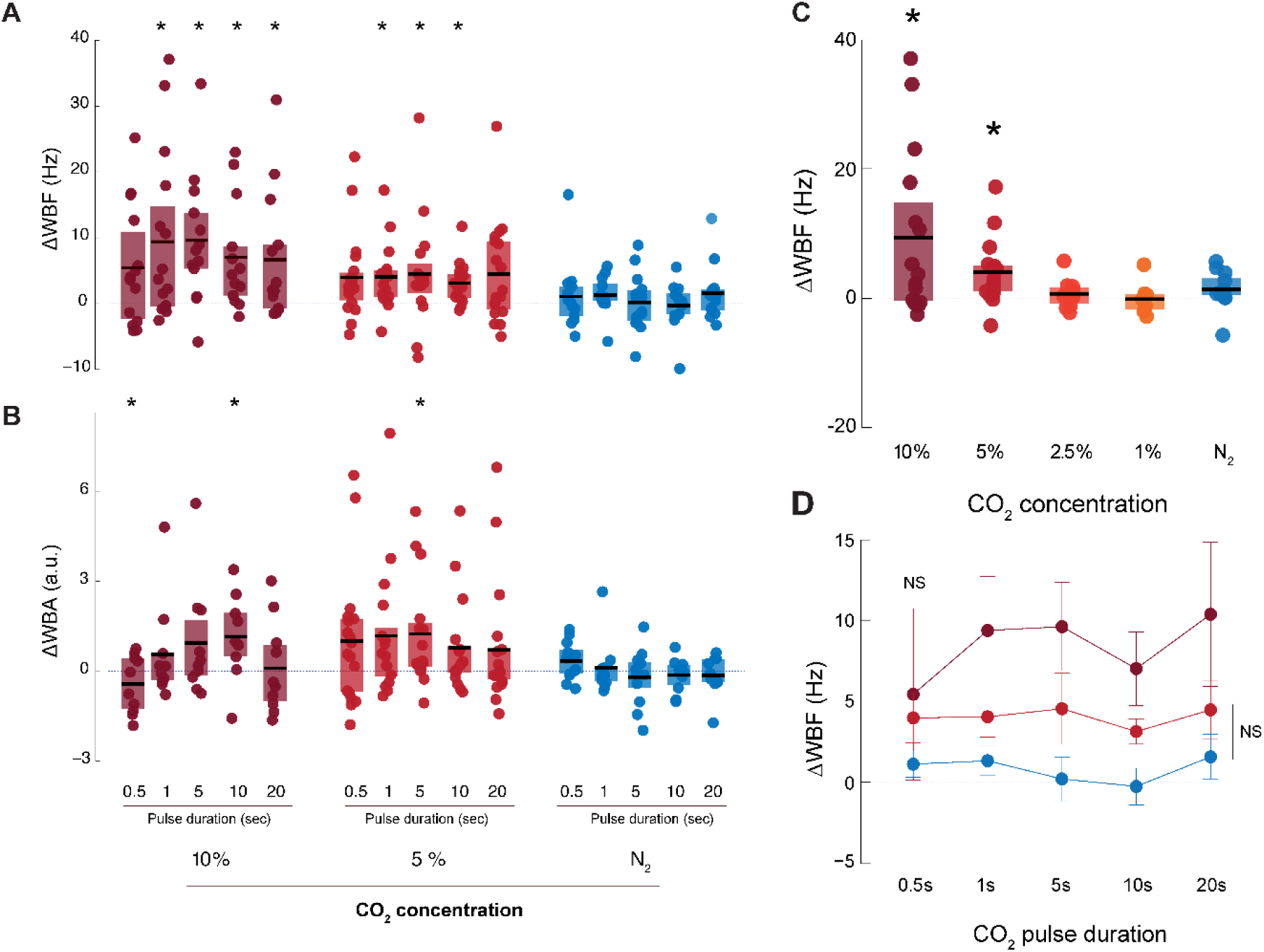
**(A)** Change in wingbeat frequency in the LED arena, after stimulation by CO_2_ at different concentrations (10%, 5%, No-CO_2_) and for various pulse durations (0.5-20s). Example schematics of ΔWBF and latency determination. **(B)** Change in wingbeat amplitude after stimulation by CO_2_ at different concentrations (10%, 5%, No-CO_2_) and for various pulse durations (0.5-20s). **(C)** Mean (±first and third quartiles) change in wingbeat frequency (ΔWBF) in response to pulses of CO_2_ at different concentrations (0 to 10%). Asterisks denote significantly elevated responses with respect to the N_2_ control (p<0.05; n = 15). **(D)** Mean (±first and third quartiles) change in wingbeat frequency (ΔWBF) in response to pulses of 10% (dark red), 5% (red) CO_2_ and N_2_ (blue) of different durations.

**Figure S2.**
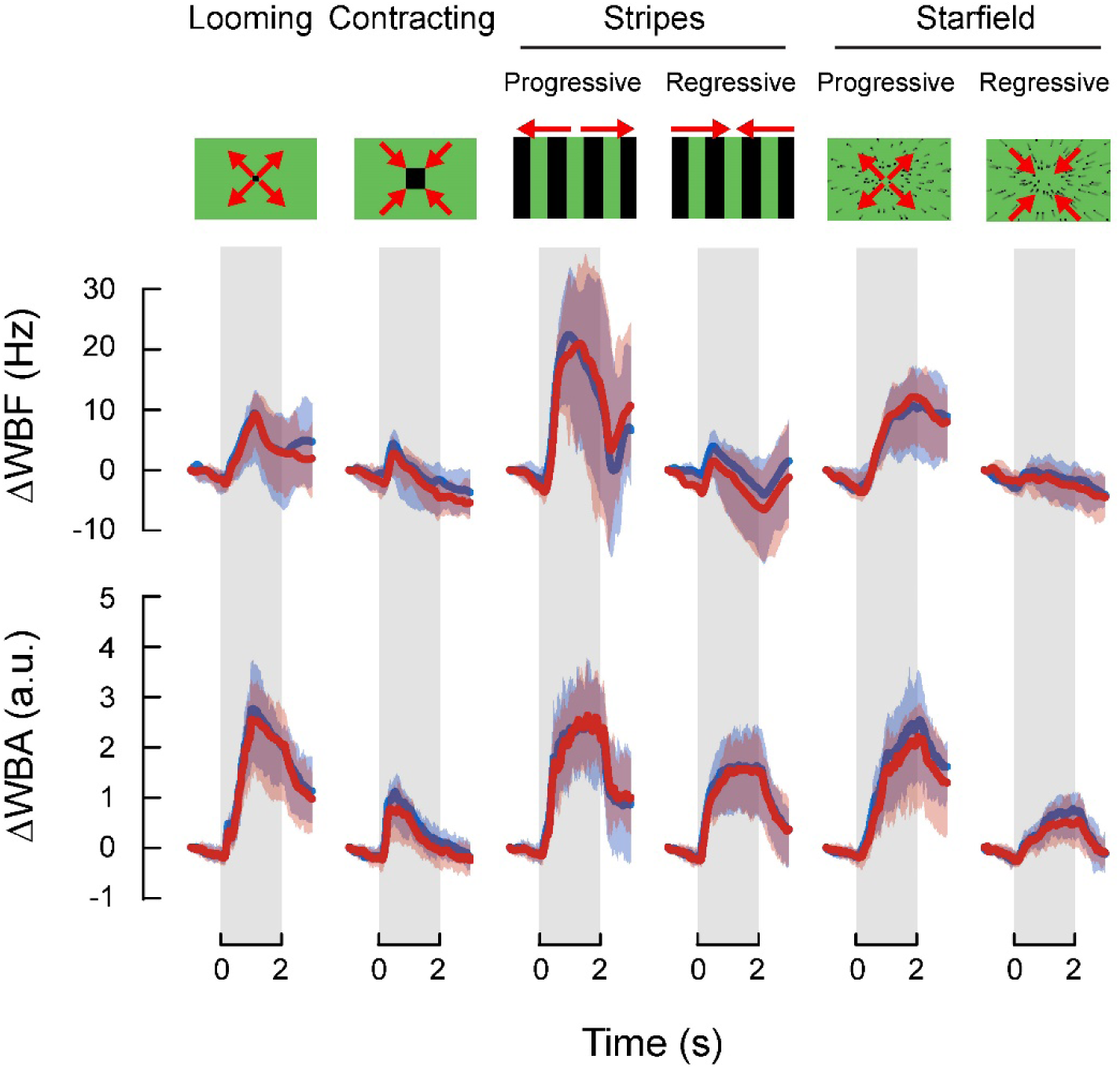
Mean responses of mosquitoes to a panel of visual stimuli. *Top*: normalized wingbeat frequency (ΔWBF), and *bottom*: amplitude (ΔWBA) changes induced by the visual stimuli. Plotted are the mean responses to visual stimuli in the absence (blue lines) and presence (red lines) of CO_2_. Shaded areas denote the first and last quartiles around the mean. Grey bars indicate the presentation of the visual stimuli.

**Figure S3.**
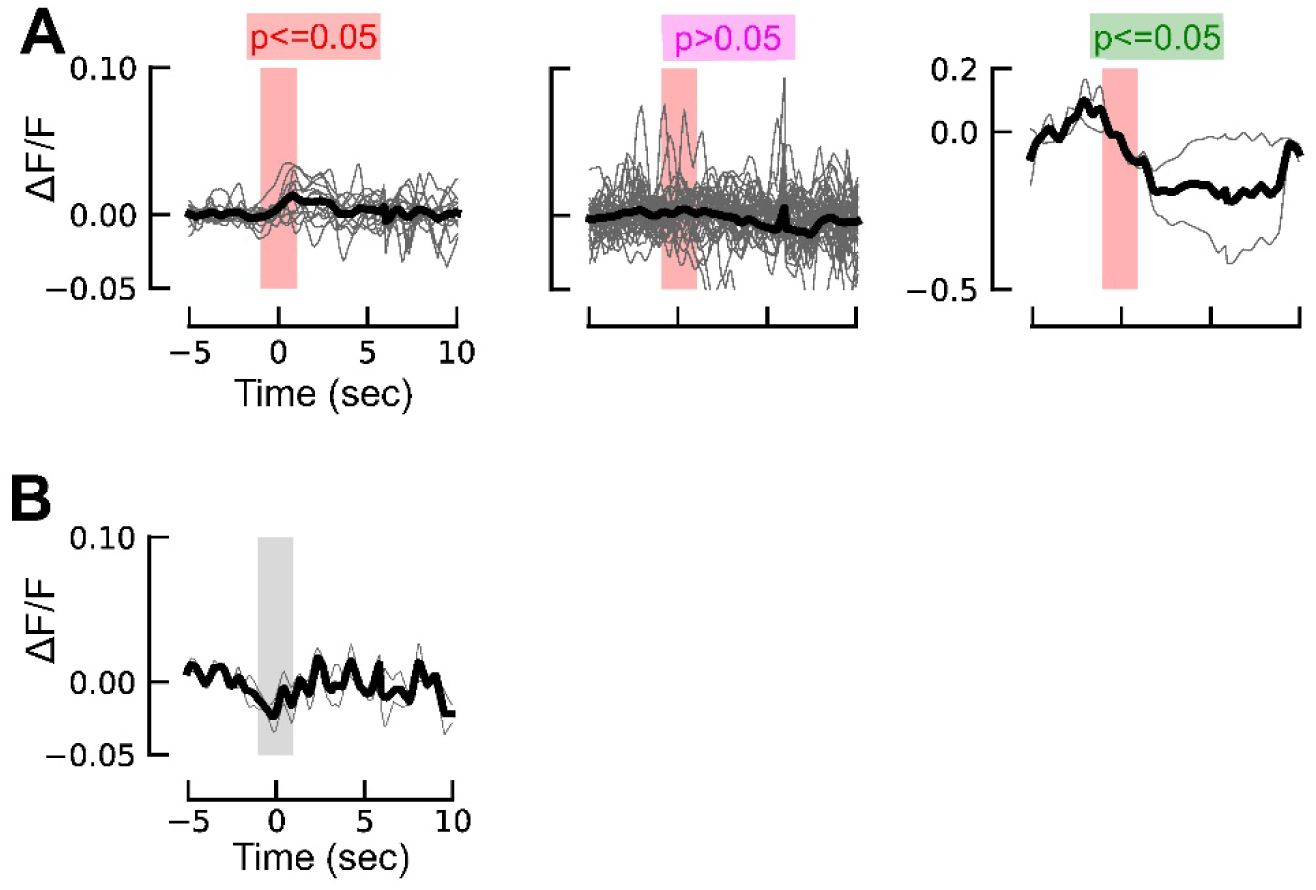
**(A)** ∆F/F time traces for odour only experiments, split into the three statistical groups shown in Figure 4E. Lobula ROIs that showed significant modulation to the odour+visual stimulus also showed significant dynamics to the odour stimulus. **(B)** ∆F/F time traces for lobula ROIs that were stimulated with pulses of elevated air flow (grey bar), showing no significant response to mechanosensory stimuli (p>0.05).

## Supplementary Tables

**Table S1.**
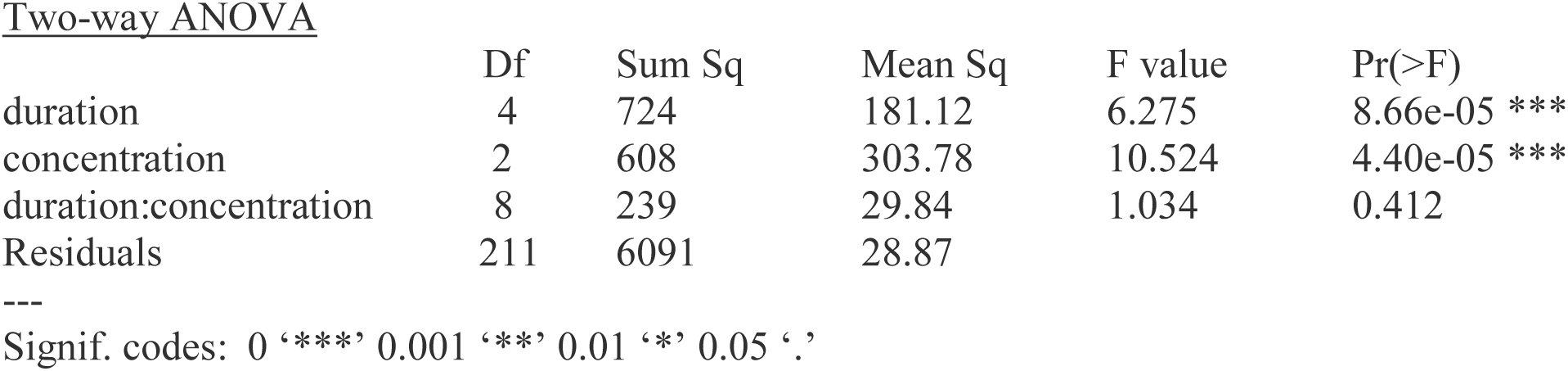
Responses to CO_2_ pulses are influenced by pulse duration and concentration.

Two-way ANOVA
|  | Df | Sum Sq | Mean Sq | F value | Pr(>F) |
| --- | --- | --- | --- | --- | --- |
| duration | 4 | 724 | 181.12 | 6.275 | 8.66e-05 *** |
| concentration | 2 | 608 | 303.78 | 10.524 | 4.40e-05 *** |
| duration:concentration | 8 | 239 | 29.84 | 1.034 | 0.412 |
| Residuals | 211 | 6091 | 28.87 |  |  |

Signif. codes: 0 '\*\*\*' 0.001 '\*\*' 0.01 '\*' 0.05 '.'

**Table S2.** Responses to visual objects are influenced by the size of the object, the lag between CO_2_ and visual stimuli, and the interaction between the two factors.

Two-way ANOVA
|  | Df | Sum Sq | Mean Sq | F value | Pr(>F) |
| --- | --- | --- | --- | --- | --- |
| Interval | 3 | 4515 | 1505 | 4.527 | 0.00403 ** |
| Square size | 3 | 65795 | 21932 | 65.973 | < 2e-16 *** |
| Interval:square size | 9 | 1326 | 147 | 0.443 | 0.91076 |
| Residuals | 294 | 97734 | 332 |  |  |

Signif. codes: 0 '\*\*\*' 0.001 '\*\*' 0.01 '\*' 0.05 '.'

## Experimental Model and Subject Details

Wild type *Aedes aegypti* mosquitoes (line Rockefeller F25, MR4-735) were used for the tethered flight experiments. The colony was maintained in a climatic chamber at 25±1°C, 60±10% relative humidity (RH) and under a 12-12h light-dark cycle. An artificial feeder (D.E. Lillie Glassblowers, Atlanta, Georgia; 2.5 cm internal diameter) supplied with heparinized bovine blood (Lampire Biological Laboratories, Pipersville, PA, USA) placed on the top of the cage and heated at 37°C using a water-bath circulation, allowed us to feed mosquitoes on weekdays. Cotton balls soaked with 10% sucrose were continuously provided to the mosquitoes. Groups of 200 larvae were placed in 26×35×4cm covered pans containing tap water and were fed on fish food (Hikari Tropic 382 First Bites - Petco, San Diego, CA, USA). Groups of 120 pupae were then isolated in 16 Oz containers (Mosquito Breeder Jar, Bioquip Products, Rancho Dominguez, CA, USA) until emergence. Adults were then transferred into mating cages (BioQuip Products, Rancho Dominguez, CA, USA) and maintained on 10% sucrose.

Mosquitoes used in the calcium imaging experiments were from of the *Ae. aegypti* Liverpool strain, which was the source strain for the reference genome sequence. Briefly, this mosquito line was generated by injecting a construct that included the GCaMP6s plasmid (ID# 106868) cloned into the piggyBac plasmid pBac-3xP3-dsRed and using *Ae. aegypti* polyubiquitin (*PUb*) promoter fragment. Mosquito pre-blastoderm stage embryos were injected with a mixture of the GCaMP6s plasmid described above (200ng/ul) and a source of piggyBac transposase (phsp-Pbac, (200ng/ul)). Injected embryos were hatched in deoxygenated water and surviving adults were placed into cages and screened for expected fluorescent markers. Mosquitoes were backcrossed for five generations to our wild-type stock, and subsequently screened and selected for at least 20 generations to obtain a near homozygous line. The location and orientation of the insertion site was confirmed by PCR (see [37] for details).

For all the experiments, 6-8 day old female mosquitoes were used. For behavioural experiments, this gave mosquitoes the time to mate in the containers before the tethered flight experiments (random dissection of females revealed that 95% of them had oocytes); all experiments in the flight arena occurred during the last three hours of the mosquitoes’ subjective day [58,59]. Female mosquitoes used in calcium imaging experiments were unmated and kept in isolation.

## Method Details

### Tethered Flight Visual Arena

Tethered flight responses by mosquitoes to olfactory and visual stimuli were tested in an LED-based arena (*sensu* [14]; Fig 1A). The arena consists of an array of 96×16 LEDs, each subtending 3.75° on the eye, subtending 360° horizontally and 60° vertically. Mosquitoes were cold anesthetized on ice and tethered to a tungsten wire using UV-activated glue (Loctite 3104 Light Cure Adhesive, Loctite, Düsseldorf, Germany) applied on the thorax. The main body axis was positioned at a 30° angle from the tether. Mosquitoes were then stored at room temperature in a closed container for an approximate 30 minute recovery period. Tethered mosquitoes were centered in a hovering position within the arena (Fig. 1A; [14]).

Mosquitoes were placed directly under an infrared (IR) diode and situated above an optical sensor coupled to a wingbeat analyzer (JFI Electronics, University of Chicago; Götz, 1987; [14,60]). The beating wings cast a shadow onto the sensor, allowing the analyzer to track the motion of both wings and measure the amplitude and frequency of each wing stroke. Measurements were sampled at 5 kHz and acquired with a National Instrument Acquisition board (BNC −2090A, National Instruments, Austin, Texas, USA).

### Odour delivery

The mosquito was centered between an air inlet and a vacuum line aligned diagonally with one another, 30° from the vertical axis (Fig 1A). The air inlet was positioned 12 mm in front of and slightly above the mosquito’s head, targeting the antennae from an angle of 15°. The vacuum line was positioned behind the mosquito 25 mm away from the tip of the abdomen. Two different airlines independently controlled by a solenoid valve (The Lee Company, Essex, CT, USA, LHDA0533115H) intersected this main air inlet, one delivering nitrogen and, the other, CO_2_. Mass flow controllers for both the CO_2_ and nitrogen delivery allowed for the CO_2_ to be set at different concentrations (0, 1, 2.5, 5 and 10%) and pulse durations. Nonanal was diluted at 1:100 in mineral oil and 2 μL was pipetted on to a filter paper (2M Whatman) in a Pasteur pipette.

### Mosquitoes responses to different CO_2_ concentrations and pulse durations

For these experiments, a visual pattern of alternating vertical stripes comprised of either inactive or fully-lit LEDs, each 16×6 pixels in size (i.e. 22.5° wide, 60° tall) was used. The pattern was briefly placed in closed loop at the beginning of the experiment in order to encourage the mosquitoes to fly and then held motionless during the presentation of CO_2_. Closed loop control of the pattern position was achieved using the difference between the left and right amplitude signals. Concentrations of 5% and 10% CO_2_ were initially tested, delivered for durations of 20, 10, 5, 1, and 0.5 seconds. One second pulses of CO_2_ at 2.5% and 1% were also tested. Potential mechanical stimulation associated with the onset of the pulses was controlled for by delivering N_2_ pulses for all the tested durations. Because a 1 sec pulse of 5% CO2 was sufficient to produce a reliable, robust frequency response, this was the concentration and pulse duration used throughout the remainder of this study.

### Moving visual patterns

To test the response to looming and drifting objects, large-field patterns of optic flow, and rotating field patterns, we adapted a broad panel of visual stimuli that are known to be important for guidance and stability during flight in other insects [30]: looming and fading squares, progressive and regressive stripes, and starfield patterns (75% of pixels ON), yaw, a square or a bar moving either from left to right or from right to left (Figs. 1, S2). The stimuli were each presented for two seconds and were separated by a 4 sec period during which all LEDs in the arena were lit. The angular velocity of objects moving on the display was 150°/sec. The entire experiment consisted of five trials of twelve visual stimuli presented twice (either immediately following a 1 sec pulse of CO_2_, or alone), the order of which was randomized at the beginning of each trial, using Matlab’s random number generator.

### Influence of odour stimulus history on visual responses

To determine how long prior, or subsequent, stimulation with an odour modifies visual responses, mosquitoes were presented with thirty-two stimuli comprised of a visual stimulus of various sizes (3.75°, 7.5°, 15°, 22.5°), moving from the left to right or right to left (yaw), and centred on the horizon of the arena. Stimuli were maintained at a constant angular velocity of 150°/sec for one revolution across a stable starfield background (75% of pixels ON). Visual stimuli were preceded by either no CO_2_, or a one second long pulse of CO_2_ delivered 1, 5 or 10 seconds before the onset of the visual stimulus (Fig. 2A). Between stimuli there was a 20 sec delay, during which the stable starfield background remained, and stimuli were presented three times to each individual. To examine whether prior stimulation of the visual stimulus modified odour evoked responses, visual stimuli were presented in the absence of odour stimuli, followed by the visual stimulus and an odour pulse occurring 1 sec after the onset of the visual stimulus. Between stimuli there was a 20 sec delay.

### Calcium imaging

Visual and odor-evoked responses were imaged in the lobula region of the mosquito optic lobe, and the antennal lobe region, taking advantage of our genetically-encoded ubiquitin-GCaMPs mosquito line [37](Figs. 3A-C, 4H). Calcium-evoked responses were imaged using the Prairie Ultima IV multiphoton microscope (Prairie Technologies) and Ti-Sapphire laser (Chameleon Ultra; Coherent). The laser power was adjusted to 20mW at the rear aperture of the objective lens (Nikon NIR Apo, 40X water immersion lens, 0.8 NA), and bandpass filtered the GCaMP fluorescence with a HQ 525/50 m-2p emission filter (Chroma Technologies) and collect the photons using a multialkali photomultiplier tube. Images were collected at 2 Hz for each visual and visual+odour stimulus, for a total duration of 350 s (Fig. 3), and calcium-evoked responses are calculated as the change in fluorescence and time-stamped and synced with the stimuli. Individual mosquitoes were tethered to a holder [16] at the center of a semi-cylindrical visual arena (frosted mylar, 20 cm diameter, 20 cm high); a video projector (Acer K132 WXGA DLP LED Projector, 600 Lumens) positioned in front of the arena projected the visual stimuli. To separate the wavelength of the light emitted by the projector from the GCaMP6 fluorescence, we used the projector’s blue channel (peak at 451 nm, 18 lux, 0.02 W/m^2^) and further reduced the longer wavelength component by covering the projector with three layers of blue gel filter (ROSCOLUX #59 Indigo). Select visual stimuli were the same as those used in the arena experiments: a stripe, square (15°) and star-field pattern (comprising 75% of the screen).

Lobula ROIs were manually selected based on three criteria, including clear expression of *PUb-GCaMP6s* (1.02 to 15.9-times the baseline fluorescence) allowing the imaging of neuropil and axons (Fig. 3), responses in at least one of the ROIs in the imaging plane to visual or olfactory stimuli (Fig. 4), and a surface area of 40-100 μm^2^. Axons and neuropil regions of interest in the lobula were imaged at approximately 40 to 100 μm from the ventral surface (Fig. 3) – neuropil in this region showed strong responses to visual stimuli and odour-evoked modulation –, and optical sections (1 μm) were taken to reconstruct neurons associated with the regions of interest (Amira v.5, Thermo Fisher Scientific). We typically have stable imaging for approximately 1.5 h allowing complete testing of the experimental series.

Antennal lobe ROIs were largely selected based on the criteria listed above, except ROI selection was based on the clear delineation between glomerular boundaries. Glomerular ROIs were imaged at 40 μm from the ventral surface, as glomeruli at this depth show strong responses to host-related odorants, including CO_2_, nonanal, octanal, and hexanoic acid, and at this depth 14-18 glomeruli can be neuroanatomically identified and registered between preparations. Moreover, GCaMP6s expression is very high in AL projection neurons (PNs), such that during odour stimulation the PNs can be imaged and 3D reconstructions can be take place *via* simultaneous optical sections with odour stimulation. Calcium-evoked responses are calculated as the change in fluorescence and time-stamped and synced with the stimulus pulses. After an experiment, the AL was serially scanned at 1 μm depths from the ventral to dorsal surface to provide glomerular registration to our tentative AL atlas (n = 6 female mosquitoes) as well as one that was previously published [60]. We note that glomeruli identified in our imaging experiments did not always conform, in terms of glomerular number and position, to the previously published atlas [60], however, the two atlases provide a first principles approach for identifying and registering glomeruli.

## Quantification and Statistical Analysis

Analyses were performed in R. For each stimulus, a baseline wingbeat frequency was determined by averaging the frequency across a 1 sec time window preceding the stimulus delivery (either visual or olfactory, according to the experiment) and then subtracting this value from the max frequency values following the stimulus. Trials were discarded in which the mosquitoes stopped flying, indicated by a drop in wingbeat frequency below 200 Hz. The mean response for each individual was calculated from the saved trials and used as a replicate to calculate the mean response for each treatment group. This latter was calculated using the difference in frequency, turning tendency (left-right amplitudes), and total amplitude (left+right amplitudes) before and after the stimulus. One-tailed Student’s t-tests for paired samples were used to test for differences from baseline and t-tests for independent samples were used to test for differences between groups. As stimuli that were presented with two directions of movement (i.e. square, stripe and yaw from left to right or from right to left) did not elicit significantly different responses (Student *t* test; 0.06<t>1.15; 40<df<84; p>0.21 for all comparisons), both directions of rotations were combined for the analysis. ANOVA and Tukey post-hoc tests were employed for multiple comparisons. When specified, multiple pairwise *t*-tests with Holm corrections were used to compare responses to visual stimuli. The delay before return to baseline wingbeat frequency was determined by determining the time at which the frequency signal crossed a threshold set at ½ standard deviation above the baseline mean frequency. The correlation between the turning response of the mosquitoes and the position of moving visual objects (i.e. squares and stripes), were quantified by means of fitted linear model defined as: mosquito.turning = α × object.position + β × object.velocity. Adjusted R^2^ values were compared by means of *t-*tests.

Calcium imaging data were extracted in Fiji/ImageJ and analyzed in python. The trigger-averaged ΔF/F were used for comparing responses to visual and odour stimuli. To statistically determine visual-evoked responses, for each ROI we assessed the difference in mean fluorescence between the time period during the visual stimulus presentation and the time period preceding the visual stimulus by comparing the two datasets to a null distribution of 10,000 bootstrapped pairwise differences drawn from the combined pre-visual stimulus and visual stimulus datasets. Similarly, for examining odour-evoked modulation, for each ROI we assessed the difference between the odour and no-odour responses by comparing the difference in the mean fluorescence during the visual stimulus for these two experiments to a null distribution of 10,000 bootstrapped pairwise differences drawn from the combined odour and no-odour datasets.

